# The evolutionary forces shaping cis and trans regulation of gene expression within a population of outcrossing plants

**DOI:** 10.1101/763870

**Authors:** Emily B. Josephs, Young Wha Lee, Corlett W. Wood, Daniel J. Schoen, Stephen I. Wright, John R. Stinchcombe

## Abstract

Understanding the persistence of genetic variation within populations has long been a goal of evolutionary biology. One promising route towards achieving this goal is using population genetic approaches to describe how selection acts on the loci associated with trait variation. Gene expression provides a model trait for addressing the challenge of the maintenance of variation because it can be measured genome-wide without information about how gene expression affects traits. Previous work has shown that loci affecting the expression of nearby genes (local or cis-eQTLs) are under negative selection, but we lack a clear understanding of the selective forces acting on variants that affect the expression of genes in trans. Here, we identify loci that affect gene expression in trans using genomic and transcriptomic data from one population of the obligately outcrossing plant, *Capsella grandiflora*. The allele frequencies of trans-eQTLs are consistent with stronger negative selection acting on trans-eQTLs than cis-eQTLs, and even more strongly on trans-eQTLs associated with the expression of multiple genes. However, despite this general pattern, we still observe the presence of a trans-eQTL at intermediate frequency that affects the expression of a large number of genes in the same coexpression module. Overall, our work highlights the different selective pressures shaping variation in cis- and trans- gene regulation.

## Introduction

Understanding why genetic variation persists in populations has long been a goal of evolutionary biology [1]. Variation within populations may be 1) neutral and maintained by mutation-drift balance, 2) deleterious and maintained by mutation-selection balance, or 3) conditionally beneficial and maintained by balancing selection [2]. The availability of large genomic and phenotypic datasets offers the potential to evaluate the relative importance of these three hypotheses by identifying the genetic loci that are associated with a trait and using population genetic approaches to determine how selection acts on these loci [3,4]. In particular, the allele frequencies of eQTLs can provide information about selection, since negative selection against deleterious mutations is expected to keep alleles at lower frequencies than they will be under neutrality or balancing selection.

Gene expression has emerged as a powerful model trait for addressing the challenge of the maintenance of variation [5]. Gene expression is a crucial aspect of the genotype to phenotype map and expression studies provide a large set of traits that can be easily measured without prior information about how these traits might relate to fitness [6]. Examining a large set of gene expression traits can reveal the evolutionary forces acting on traits in general, rather than a few or a handful of pre-defined traits chosen for specific reasons. The genetic variation that shapes expression can be partitioned into two categories: cis-regulatory variants that only affect the allele they are linked to and trans-regulatory variants, that affect both alleles equally and can be located near or far from the gene they regulate [7,8]. Previous work has mapped the genetic variants that affect expression (eQTLs) of nearby genes and shown that local eQTLs and cis-eQTLs are generally under negative selection [9–12]. However, trans-eQTLs may be under different selection pressures than cis-eQTLs. Since trans-regulatory variation can affecs the expression of multiple genes, trans-regulatory elements may have greater pleiotropic effects on phenotypes and be subject to stronger negative selection than cis-regulatory variants [13]. This prediction is supported by evidence of greater trans-regulatory variation within species compared to between species [7,14], reduced population frequencies of distant eQTLs compared to local eQTLs [15], and greater effect sizes of standing cis-regulatory variation than trans regulation [5,16,17], although these effect size differences may also be caused by differences in mutational input [18].

Despite the expectation that purifying selection will reduce trans-acting regulatory variation within species, there is evidence that trans-regulatory variation is common. Linkage mapping from crossing experiments and population-based association mapping have often found trans-regulatory hotspots, where genetic variation at a locus affects expression of numerous genes [11,16,19–23] (but see [17]). Segregating trans-variation is more likely to be tissue-specific than cis-regulatory variation in humans [24] and, in *Arabidopsis thaliana*, trans-eQTLs are particularly important for expression changes in response to drought [23,25]. These findings suggest that trans-eQTLs contribute to standing variation, especially in specific tissues and environments.

Here we both map trans eQTLs for single genes and look for loci associated with the expression of many genes [26–28], in a single population of the plant *Capsella grandiflora*, an obligately outcrossing member of the Brassicaceae family with large effective population size and high levels of genetic sequence diversity [29,30]. To look for eQTLs affecting the expression of multiple genes, we use coexpression networks to summarize expression across many genes and test for associations between genetic variants and the expression of network modules. Coexpression networks are a powerful way to find patterns in large transcriptomic datasets [17,31–35]. For example, coexpression networks made across conditions, tissues, and developmental time can successfully identify specialized metabolic pathways [32] and coexpression modules made with a diverse panel of mouse lines correlate with phenotype [36]. In addition, changes in coexpression module expression have been linked to adaptation [37] and changing ecological conditions [35]. We detect a large number of putative cis and trans eQTLs and show that, based on allele frequencies, trans eQTLs are generally under stronger negative selection than cis eQTLs. We use coexpression networks to summarize expression levels across many genes and detect four eQTL for coexpression module expression. Overall, our results suggest that negative selection acts on trans eQTLs more strongly than cis, but there are some trans-eQTLs affecting large numbers of genes at appreciable frequencies in the population.

## Results

### Linking allele frequencies and selection in cis and trans eQTLs

We tested for associations between leaf expression at all genes with 1,873,867 tag SNPs in 146 individuals. We identified 6,231 associations (FDR < 0.1) between 5,468 unique snps and 2,341 genes **(Figure 1)**. These eQTLs were associated with the expression of between 1 and 93 genes. We separated associations into 2,472 trans eQTLs that were more than 5kb away from the gene they regulated and 3759 local (putatively cis) eQTLs that were less than 5kb away from the gene they regulated. We will refer to these eQTLs as cis eQTLs for clarity, while noting that some of them may be caused by trans-eQTLs located near the genes they regulate. 3,300 of these cis-eQTLs were detected previously in a subset of 99 individuals at p<0.05, and 2,636 at an FDR of 0.1 (the cutoff used in that study) [9]. Trans-eQTLs had larger effect sizes than cis-eQTLs (p < 0.001, mean local effect size = 0.91, mean trans effect size = 1.01).

**Figure 1:**
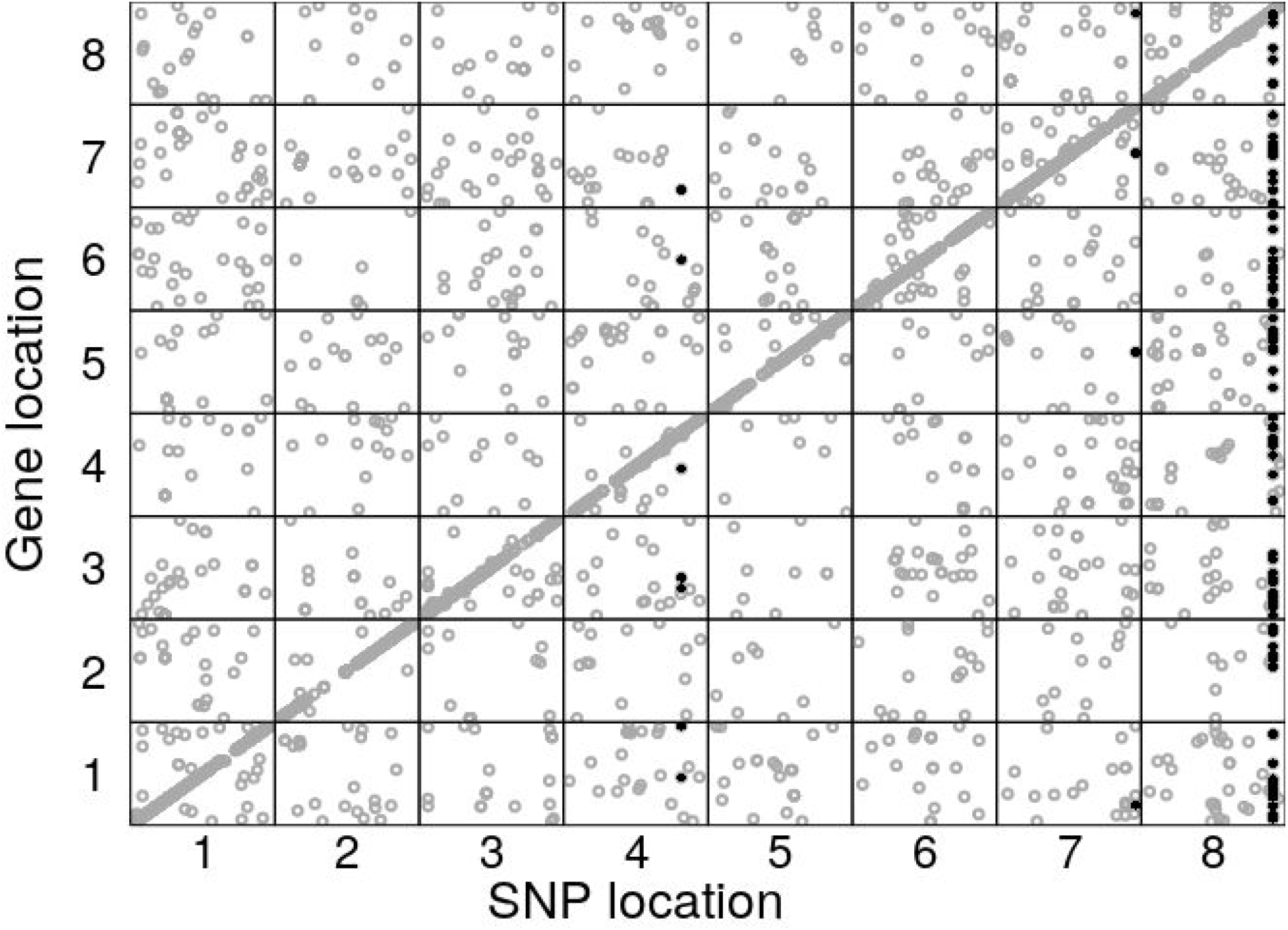
All-by-all eQTL locations. Each point represents a SNP, whose location is described on the X axis, that is associated with the expression of a gene whose location is described on the y axis (FDR < 0.1). Black points are SNPs that were also identified in the coexpression eQTL analysis.

We used the minor allele frequency (‘MAF’) to infer the relative strength of selection acting on different types of eQTLs. Trans-eQTLs had lower MAFs than than cis-eQTLs (**Figure 2**, p < 0.001, mean trans MAF = 0.214, mean cis MAF = 0.267). This result was robust to the cutoff distance used to define cis-and trans-eQTLs: trans-eQTLs had lower MAFs than cis-eQTLs for distance cutoffs of 1 kb, 2.5 kb, and 10 kb (p < 0.001). This difference in allele frequencies is consistent with stronger negative selection on trans-eQTLs than cis-eQTLs. However, since trans-eQTLs have larger effects on expression, there may be more power to detect trans-eQTLs at lower frequencies, causing the observed pattern. We found that when we restricted our analysis to eQTLs in the top quartile of effects (684 cis-eQTL and 692 trans eQTL), trans-eQTLs were still present at lower minor allele frequencies than cis-eQTLs, consistent with allele frequency differences resulting from stronger negative selection on trans-eQTLs than cis-eQTLs (p < 0.001, mean cis MAF = 0.115, mean trans MAF = 0.098).

**Figure 2.**
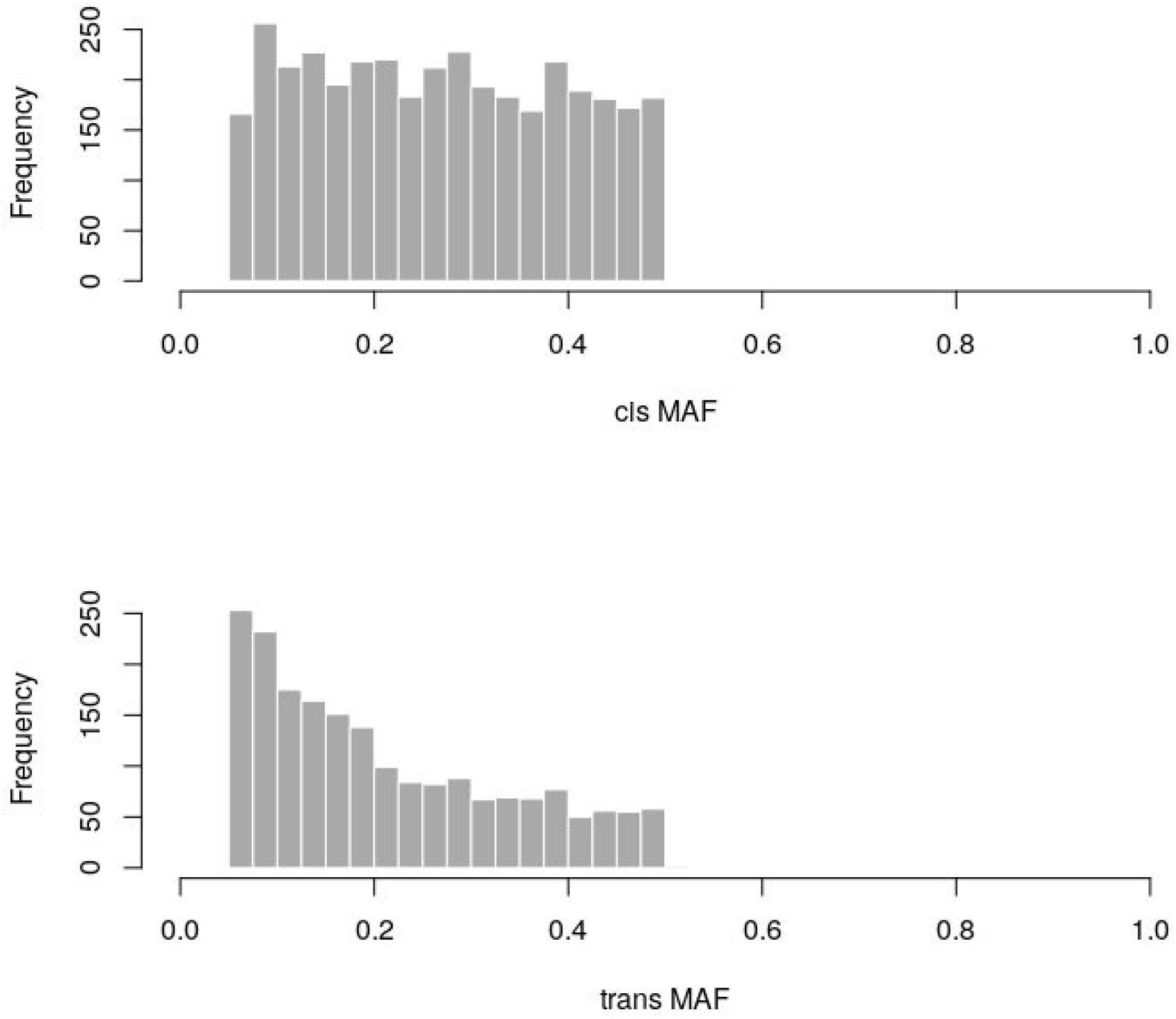
Distribution of minor allele frequencies (MAF) of cis and trans eQTLs.

We also investigated the MAFs of eQTLs associated with the expression of many genes compared to the MAFs of eQTLs associated with only one gene. We restricted this analysis to trans eQTLs, to ensure that the differences between cis and trans eQTLs described above did not confound the results. Within trans eQTLs, there were 1,635 eQTLs associated with one gene and 376 eQTLs associated with the expression of more than one gene at an FDR < 0.1. The MAF of eQTLs with one association was significantly higher than the MAF of eQTLs with more than one association (p < 0.001, one gene MAF = 0.214, more than one gene MAF = 0.196). This result is consistent with negative selection acting more strongly on eQTLs with many associations than eQTLs with one one association.

### Coexpression module GWAS identifies SNPs affecting expression modules

We identified 16 coexpression modules ranging in size from 69 to 6392 genes **(Figure S1)**. We summarized expression level across modules, which we will refer to as ‘module expression’, for each individual using eigengenes. Module expression is not correlated with collection timing **(p > 0.2, Figure S2).** Module expression values show varying distributions: some modules had normal distributions, some were bimodal, and some showed strong skews where a few individuals had very high module expression compared to other individuals **(Figure S3)**. Because the skewed distribution of module expression values could lead to false positives during association mapping, we quantile-normalized expression level for association mapping.

Genome-wide association mapping for module expression identified four SNPs associated with the expression of two modules (FDR < 0.1, **Table 1)**. We refer to these SNPs as ‘coexpression-eQTLs’. All four of the coexpression-eQTLs were also identified as eQTLs in the all-by-all analysis. Two coexpression-eQTLs are associated with expression of the ‘lightyellow’ module and two with the expression of the ‘white’ module. **(Figure 3, Table 1)**. Both eQTLs for the ‘white’ module were located near each other (1.2 kb apart). We mapped the association between all SNPs (not just tagging SNPs) and ‘white’ module expression in this region and found additional significant associations (**Figure 3B)**, suggesting that there is a longer block of loci in linkage disequilibrium associated with module expression.

**Table 1:**
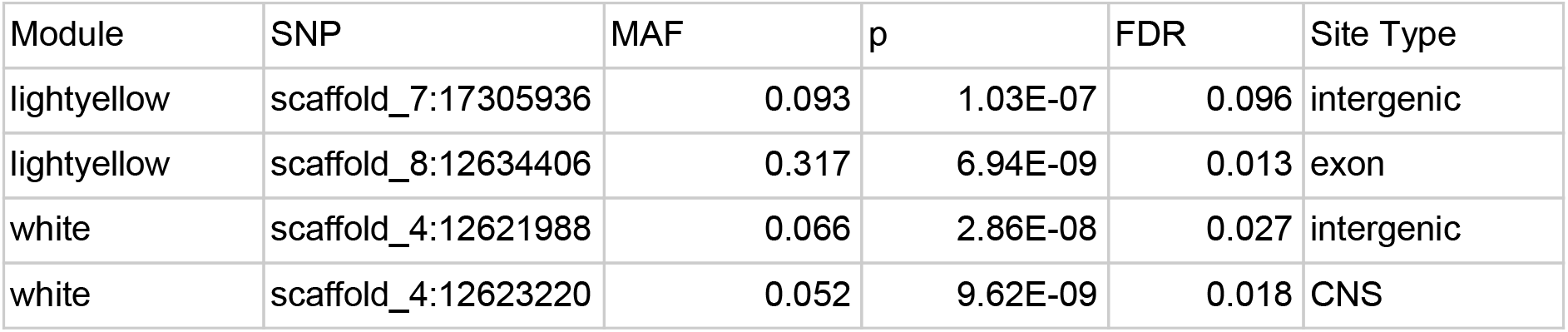
Information about significant coexpression eQTLs (FDR < 0.1). ‘CNS’ stands for ‘conserved noncoding sequence’, ‘MAF” stands for minor allele frequency

| Module | SNP | MAF | p | FDR | Site Type |
| --- | --- | --- | --- | --- | --- |
| lightyellow | scaffold_7:17305936 | 0.093 | 1.03E-07 | 0.096 | intergenic |
| lightyellow | scaffold_8:12634406 | 0.317 | 6.94E-09 | 0.013 | exon |
| white | scaffold_4:12621988 | 0.066 | 2.86E-08 | 0.027 | intergenic |
| white | scaffold_4:12623220 | 0.052 | 9.62E-09 | 0.018 | CNS |

**Figure 3:**
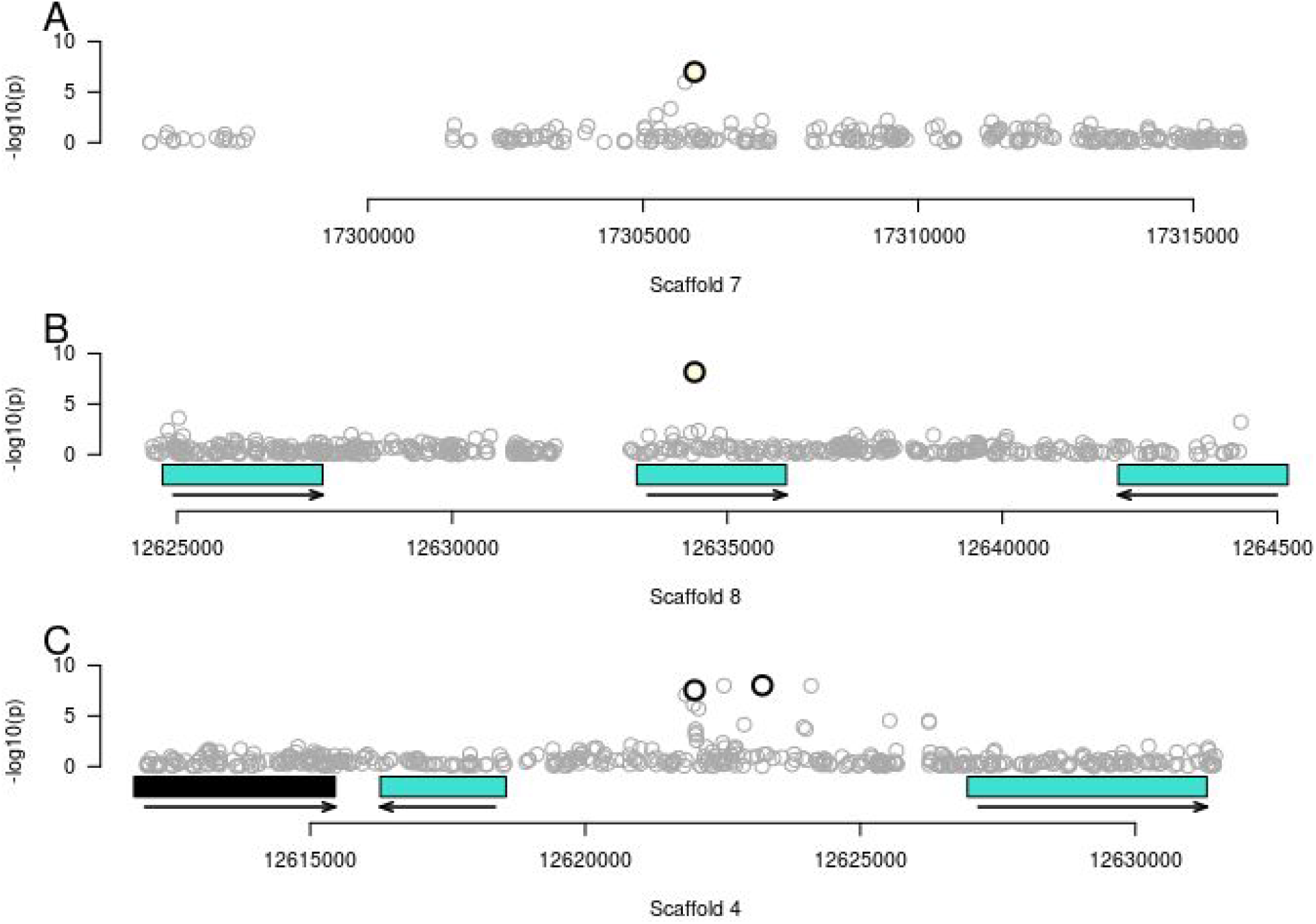
Physical locations of coexpression QTLs. Coexpression eQTLs are represented by white points with black borders. All other SNPs are plotted in gray. These plots include associations for all snps, not just the tagging snps, plotted by location, on the x axis, and the significance of association with the module indicated by color on the y axis. The locations of nearby genes are shown by rectangles, colored by the module the gene is in. The direction of transcription of each gene is shown by a black arrow. A) The first coexpression eQTL on Scaffold 7, B) the second coexpression eQTl on scaffold 8, and C) the third and fourth coexpression eQTLs on Scaffold 4.

We further investigated the gene containing the one coding coexpression eQTL, which is associated with expression of the ‘lightyellow’ module. This eQTL was in a 4-fold degenerate site of the gene Carubv10025970m. Its closest ortholog in *Arabidopsis thaliana*, AT5G65683.1 or WAV3 HOMOLOG 2 is a member of the WAVY GROWTH 3 E3 ligase family which is involved in root gravitropism but also shows expression in Arabidopsis young leaves. This coexpression eQTL is also associated with the expression of 93 genes in the all-by-all analysis. All but one of these 93 genes was in the “light yellow” module. The minor allele frequency of this eQTL is 0.317.

### Relating coexpression modules to traits

We also conducted GWAS on phenotypic traits (days to bolting, days to flower, leaf nitrogen content, leaf carbon content and leaf shape traits) following the same procedures described above for coexpression modules. No associations were significant at a FDR < 0.1 or even at an FDR < 0.25. Module expression was correlated with a number of trait measurements. There were four modules whose expression was correlated with days to bolt (**Figure 4**, p < 0.05 after Bonferroni correction for 16 tests). None of the four coexpression eQTLs detected were significantly associated with any phenotypes.

**Figure 4:**
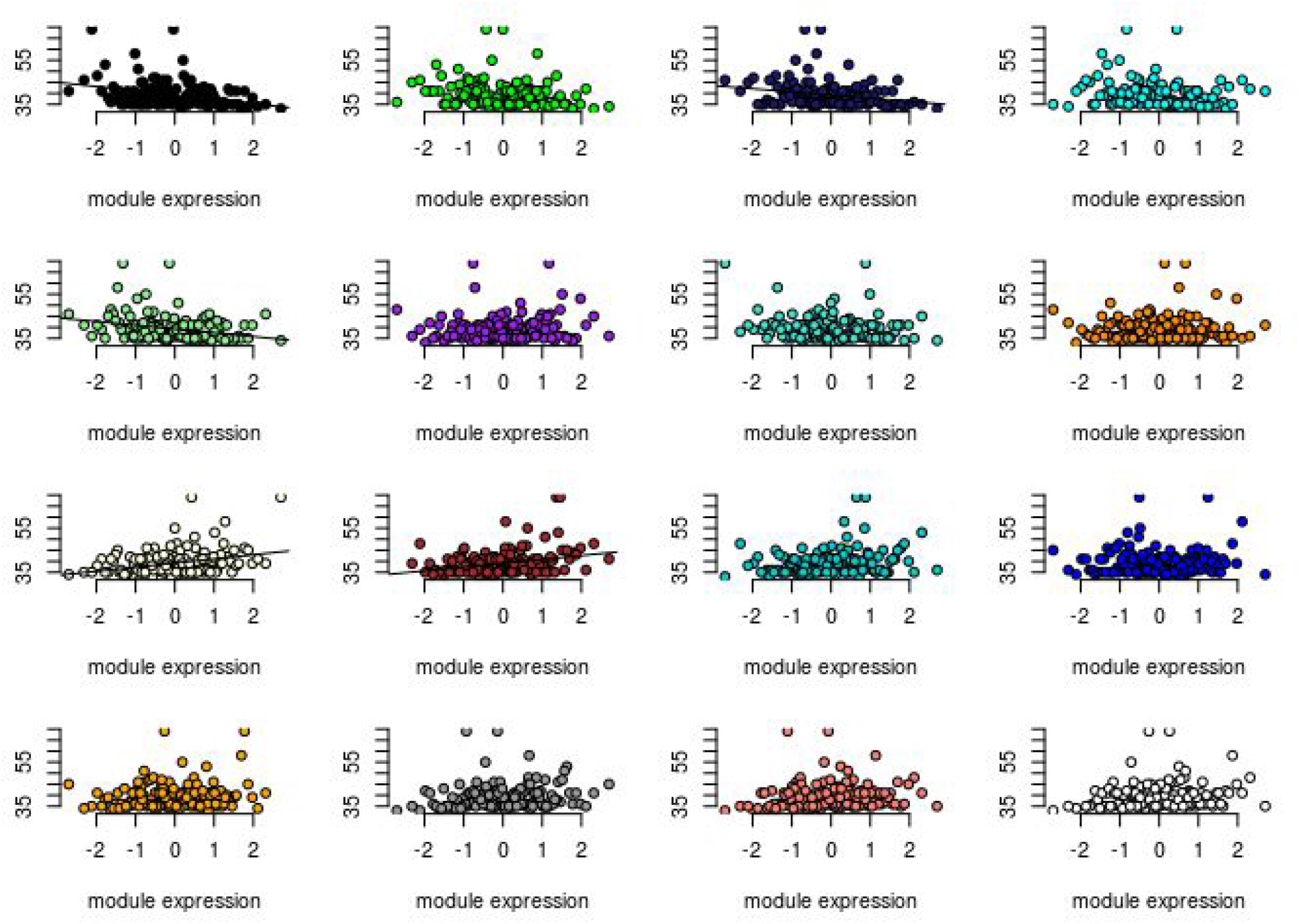
Correlations between module expression level and traits. The correlation between expression of each module (x axis) and days to flower (y axis). If the correlation was significant (p < 0.05 after Bonferonni correction), there is a black trend line drawn.

### Population genetic signatures of selection on eQTLs

We compared signatures of selection around local and trans eQTLs identified by the all-by-all analysis and found that trans-eQTLs were in windows that had lower Tajima’s D on average (Tajima’s D = −0.596) than cis-eQTLs (Tajima’s D = −0.455, p < 0.0001), consistent with the regions around trans eQTLs being under stronger negative selection than regions around cis-eQTLs, although this could also result from differences in the genomic landscapes of trans-eQTLs compared to cis-eQTLs.

While we have evidence that local cis-regulatory eQTLs are in general under negative selection in this population [9], and our present analysis suggests that all-by-all trans-eQTL are subject to stronger puriifying selection, we were curious if we could detect evidence of recent positive or balancing selection on eQTLs detected in the coexpression eQTL analysis as well as in the all-by-all analysis. We measured **π** and Tajima’s D at putatively neutral sites across the genome in 500 bp windows and used SweeD to test for evidence of selective sweeps in 50 SNP windows. None of the coexpression eQTLs were located in windows that were outliers (top 2.5% of windows) for **π**, Tajima’s D, or sweep likelihood (**Fig. S4, Fig. S5, Fig. S6**).

## Discussion

In this study, we have mapped the genetic basis of genome-wide expression variation within a single population of an outcrossing plant. The allele frequencies of trans-eQTLs suggest that the variants that affect trans-regulation are under stronger negative selection than cis-eQTLs, and that trans-eQTLs associated with the expression of multiple genes are under stronger negative selection than trans-eQTLs associated with the expression of only one gene. In addition, windows containing trans eQTLs have lower Tajima’s D than window containing cis-eQTLs, also consistent with stronger negative selection acting on trans-eQTLs. However, despite the general pattern of negative selection acting on trans-eQTLs, we detected four eQTLs associated with the expression of coexpression network modules, one of which is independently associated with the expression of 93 genes and present at an intermediate frequency. Overall, this work suggests that trans-eQTLs are under different selective pressures than cis-eQTLs, but that within trans-eQTLs there is also a great deal of variation in selection.

Our results are consistent with previous work showing that distant eQTLs are at lower minor allele frequencies than local eQTLs in *Arabidopsis thaliana[15]*. However, the *A. thaliana* result comes from association mapping done in a diverse panel of lines from across the species range where allele frequencies could be shaped by negative selection or local adaptation, so our results reflect a clearer indication that negative selection acts more strongly on trans-eQTLs than cis-eQTLs. In addition, the pattern that trans-eQTLs that are associated with the expression of multiple genes are at lower minor allele frequency than trans-eQTLs with only one association is consistent with evidence that negative selection acts more strongly on pleiotropic loci in *C. grandiflora [34]* and in other species [13,38].

One important aspect of our use of coexpression modules in the eQTL analysis is that we used “genotype networks’ generated from expression data measured in the same tissue type at the same time in a set of genetically distinct individuals. Therefore, the coexpression modules we observed were shaped by genetic perturbations, not tissue or developmental differences. While coexpression measured across multiple timepoints (‘developmental networks’) has been linked to functional relationships [39,40], coexpression modules generated from genetically distinct individuals have different properties than those generated from different tissue types [17,41]. In some cases, this difference is helpful: analyses combining GWAS and coexpression networks have the most power when using coexpression networks made from genetically distinct samples [41]. However, it is important to keep in mind that the expression datasets used will affect coexpression modules.

Mapping eQTLs has furthered our understanding of the nature of genetic variation maintained within natural populations. Analyses combining genomic and transcriptomic data from natural populations are relevant in the context of models using transcriptomic data to build a mechanistic understanding of the evolutionary forces maintaining variation within populations [42–44]. In addition, since gene expression is important for adaptive divergence [45–47], understanding the maintenance of genetic variation for expression is important for understanding how organisms will adapt to new environments.

## Materials and Methods

### Genomic, transcriptomic, and phenotypic data

All genomic and transcriptomic sequence data was previously published in Josephs et al. (2015) and Josephs et al. (2017b). We briefly describe data generation here. We collected individuals from a single population of *C. grandiflora* individuals located near Monodendri, Greece. We conducted a generation of random crosses in the greenhouse, and then grew 146 individuals descended from these random crosses in a growth chamber with 16 hours of daylight at 22°C. We measured traits on these individuals and collected RNA from leaf tissue collected 39 days after planting. Leaves were collected and flash frozen at night sequentially in 9 roughly-equally sized bins, which we will refer to as “collection bin”. We extracted RNA using Qiagen RNAeasy kits.

We extracted DNA from leaf material using a CTAB procedure. Both RNA and DNA was sequenced at the Genome Quebec facility with Hiseq 2000 with Truseq libraries with 100bp long reads. DNA was mapped to the standard *C. rubella* reference genome [48] with Stampy [49] and RNA was mapped to an exon-only reference genome using Stampy as well. SNPs were called from the genomic sequence data using GATK Unified Genotyper [50] and expression levels were measured with HTseq and normalized for sequencing depth by dividing by the median expression level for each individual. [51]. We did not detect interactions between GC content, expression level, and lane [9]. We used the ComBat function from the sva R package to adjust expression level for collection batch [52].

In addition to collecting RNA and DNA for sequencing on these 146 individuals, we measured a number of phenotypes. We measured days to bolting and days to flowering daily (measured since planting date). We collected leaves at day 49 after planting, scanned leaves, and measured leaf shape as reported in [53]. Briefly, dissection index was calculated as DI = (perimeter^2^)/(4π*area), so that a circle of the same area would have a value of 1.0 and increasing values indicate increasing complexity and alpha shape dissection index is a similar parameter, but for alpha shapes. We measured leaf carbon and nitrogen content in one leaf per individual. Leaves were collected at day 49 after planting, dried, and ground to powder for elemental analysis by the Ecosystems Analysis Lab at the University of Nebraska. We note that both shape and elemental data came from different leaves than the RNAseq data. We estimated Pearson and spearman correlations between module expression and trait values with the cor.test function in R [54].

### Building coexpression networks

We used the program WGCNA [55](version 1.68 running on R version 3.6.2) to identify coexpression modules present within the 145 transcriptomes using the expression level of all genes with median expression greater than 5 reads per gene (n = 18,806). The coexpression analysis groups together genes with similar patterns of pairwise correlation of expression. We were interested in retaining the information embodied in the sign of the gene expression correlations, so we conducted a signed network analysis. We used a soft thresholding value of 12, as suggested by the authors of the WGCNA package for signed networks and a minimum module size of 30. Genes that exhibited similar patterns of connectivity (i.e., genes showing high “topological overlap”) were grouped together in the same coexpression modules, based on hierarchical clustering of topological overlap values, in which a dynamic branch-cutting algorithm was used to define initial gene co-expression modules. Module eigengenes (the first principal component of the gene expression values of modules) were calculated, and modules whose eigengenes were correlated at a level greater than 0.8 were merged to arrive at the final set of co-expression modules. The resulting modules were labeled with different colors for ease of referencing[34]. We investigated specific eQTLs in the Joint Genome Institute’s Phytozome v12.1 genome browser for *Capsella rubella* v1.0[48] and investigated specific orthologs in *Arabidopsis thaliana* using TAIR[56].

Some of the modules had expression levels that were very skewed, such that a few individuals showed extremely high module expression compared to the rest of the individuals **(Fig S1)**. To reduce potential false-positives in the association mapping study due to skewed expression levels, we quantile normalized module expression levels using the qqnorm function in R [54].

### Association mapping

We tested for associations between SNP genotype and individual gene expression, phenotypes, and module expression. For all association mapping analyses, we filtered out SNPs with a minor allele frequency below 0.01 and more than 0.05 missing data, leaving 5,560,798 SNPs. We used Haploview to identify 1,873,867 tag SNPs with minor allele frequency > 0.05 that described the dataset.

We tested for associations between the tag SNPs with the expression of 18,806 genes using the linear model in Matrix eQTL [57]. We quantile normalized gene expression levels using the qqnorm function in R to reduce false positives caused by skewed expression distributions. While all samples came from the same population, we controlled for residual population structure by generating a centered kinship matrix with GEMMA [58] and including the first five principal components of the kinship matrix as covariates.. Since all tag SNPs were tested against all genes, we conducted 35,587,985,444 tests. Matrix eQTL estimates false discovery rates using a Benjamini–Hochberg procedure.

After detecting eQTLs, we compared cis and trans eQTLs. We defined putative cis eQTLs as eQTLs that were less than 5 kB from the transcription start or end site of the gene they were associated with and trans eQTLs as eQTLs that were either on a different chromosome from the gene they were associated with or more than 5 kB away from the transcription start or end site of the chromosome. We compared the minor allele frequencies and effect sizes of eQTLs using the t.test function in R [54].

We did association mapping with GEMMA [58] on module eigengenes (PC1 of expression values of a module), morphological, and life history traits as our phenotypes. We controlled for residual population structure using the standardized kinship matrix and SNPs with minor allele frequency > 0.05 and missing data < 0.05. We used the likelihood ratio p values [59] and calculated the p-value cutoffs corresponding to a false discovery rate of 0.1 for each trait and module expression level using the FDR method of the p.adjust function in R [60].

### Population genetic signatures of selection

We used genomic sequence from 188 individuals published in [9]. We downsampled all sites to 320 chromosomes per site and then calculated pi and Tajima’s D in 500 bp windows across the genome at non-coding (excluding conserved non-coding sites from [61]), intronic, and 4-fold degenerate sites. We used SweeD to calculate the likelihood of a selective sweep occuring on every 50th SNP (windows were ~600 bp wide on average) using non-conserved intergenic, intronic, and 4-fold degenerate sites for 182 individuals from the focal population [62].

## Acknowledgements

We thank Niroshini Epitawalage, Amanda Gorton, Robert Williamson and Jasmina Uzunović for research assistance, as well as Graham Coop, Jeff Ross-Ibarra, and members of the Stinchcombe, Wright, Coop and Ross-Ibarra labs for helpful comments and advice. This work was supported in part by Michigan State University through computational resources provided by the Institute for Cyber-Enabled Research. This work was funded by a National Science Foundation Graduate Research Fellowship (DGE-1048376) and a National Science Foundation Plant Genome Postdoctoral Fellowship (IOS- 1523733) to EBJ, Natural Sciences and Engineering Research Council of Canada Discovery Grants to DJS, SIW, and JRS and a Value-directed Evolutionary Genomics Initiative grant (Genome Quebec/Genome Canada) to DJS, SIW, and JRS.

---

**Figure S1:**
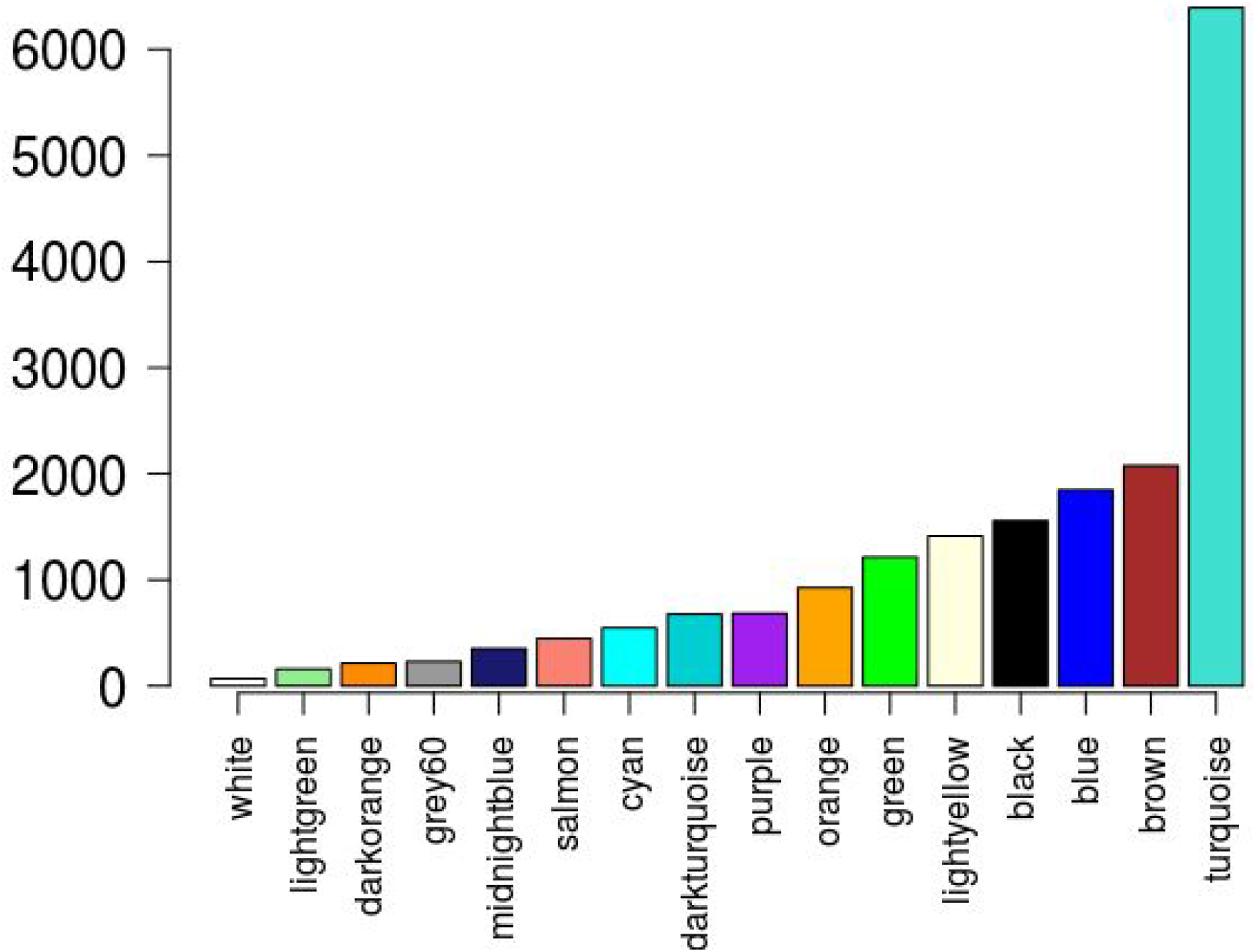
The number of genes in each module.

**Figure S2.**
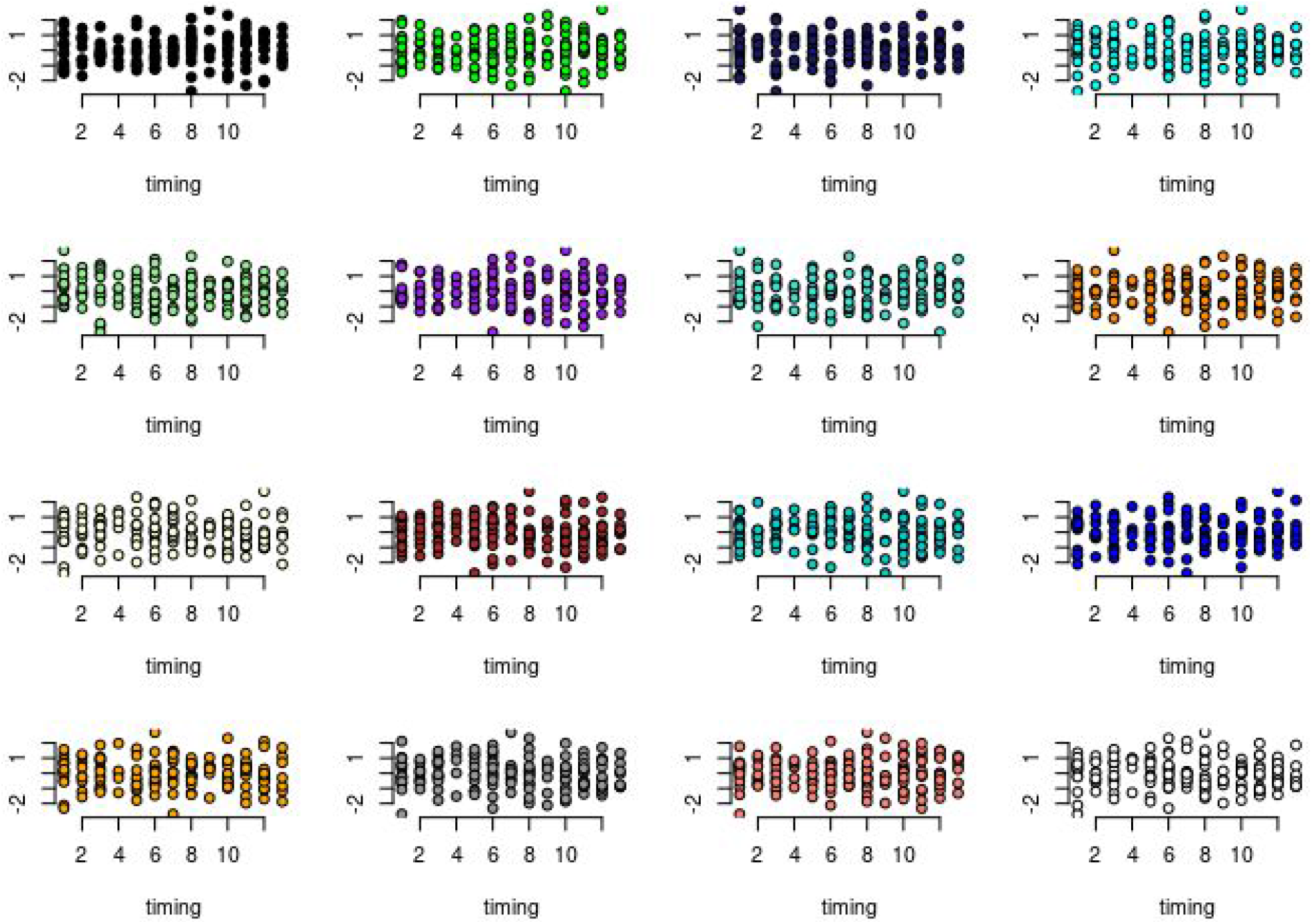
Relationship between collection box (x axis) and module expression (y axis) for 16 coexpression modules. Each point represents one plant.

**Figure S3:**
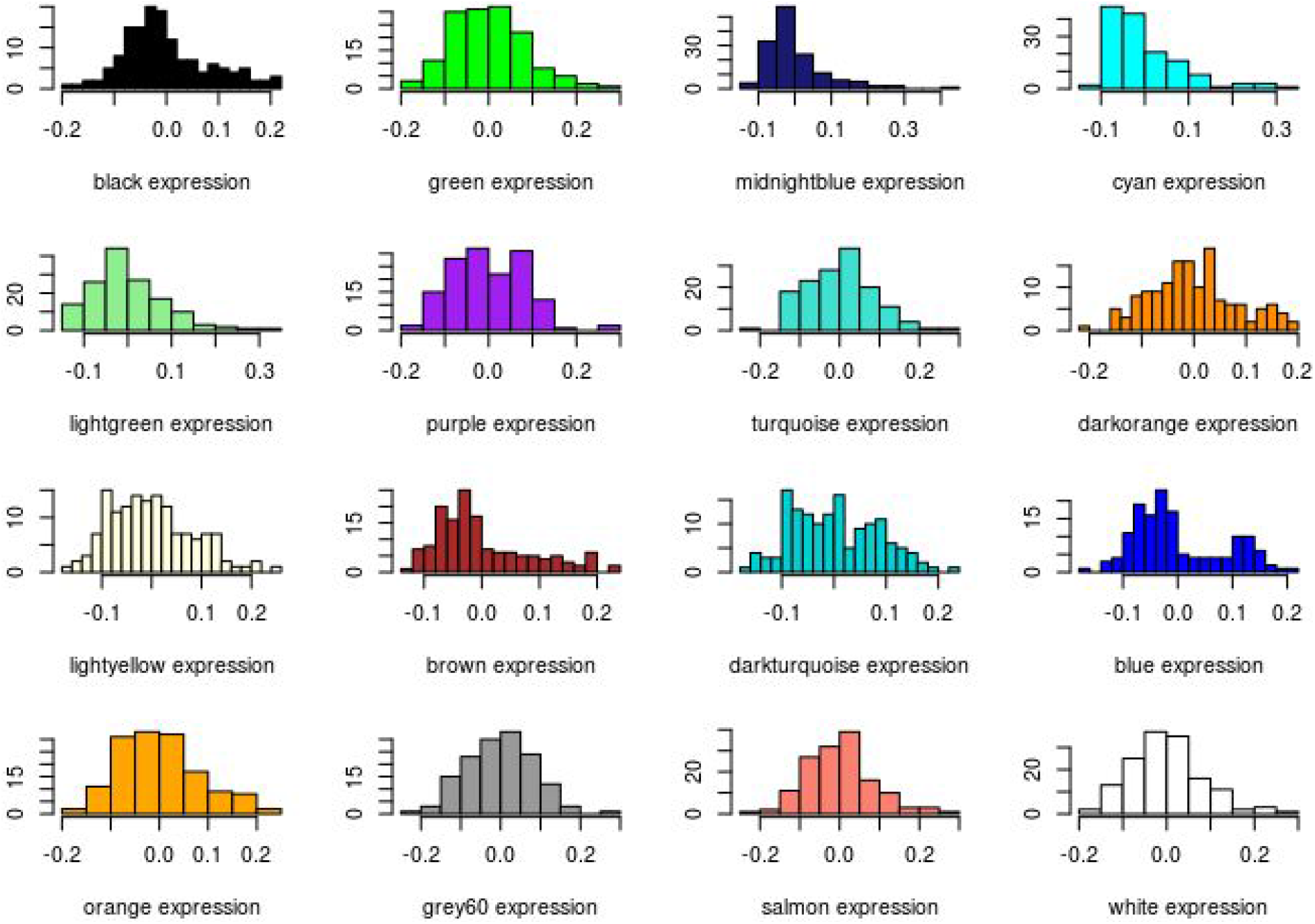
Distribution of un-normalized eigengenes. Each plot shows a histogram of eigengene expression for a specific coexpression module. The module name is labeled below the x axis.

**Figure S4:**
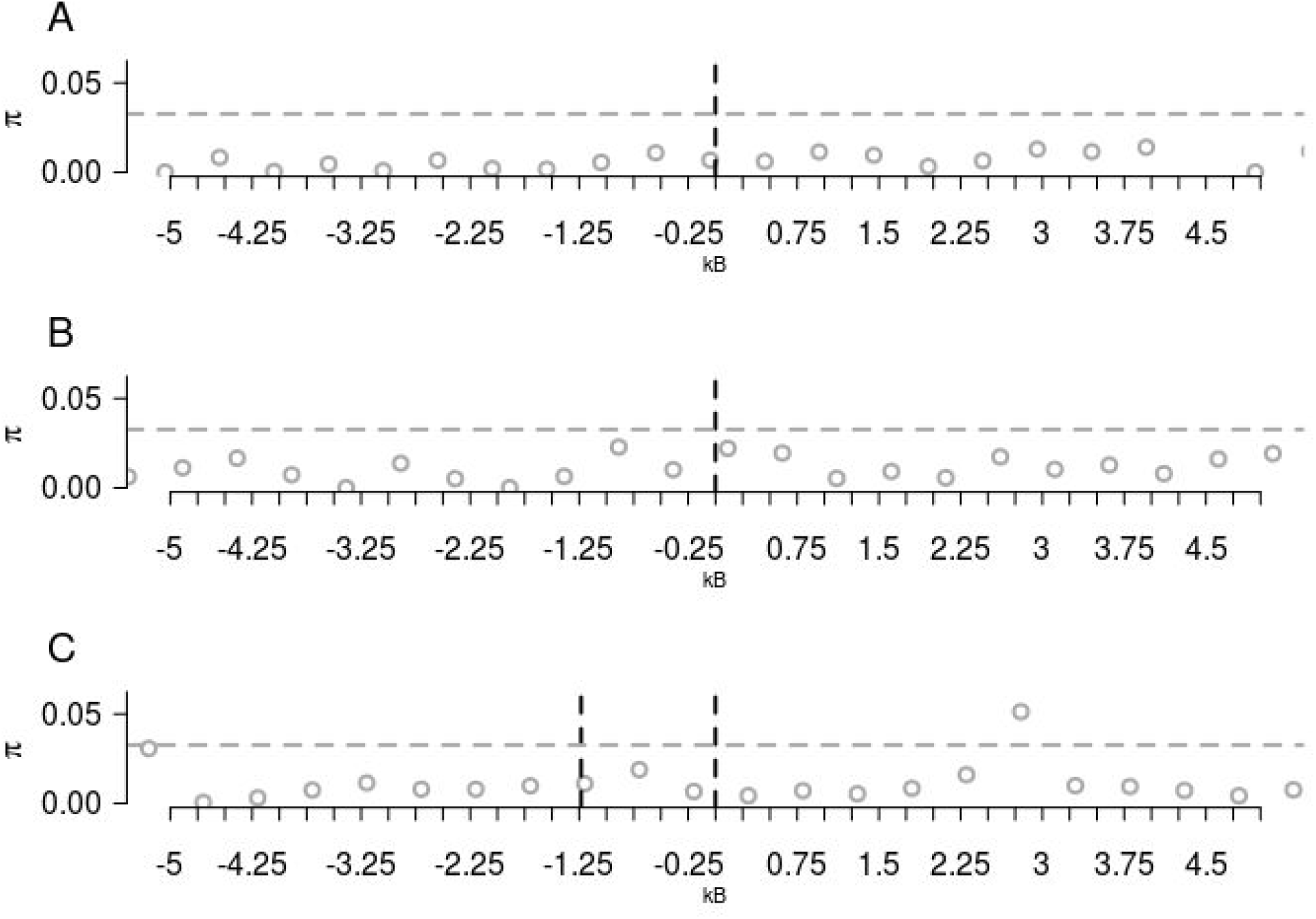
Diversity (**π**) at putatively neutral sites in 500 bp windows around coexpression-eQTLs. Gray horizontal dotted lines show 95% cutoffs of the observed distribution of likelihoods and black vertical lines show the locations of eQTLs. Each panel corresponds to the eQTL shown in **Fig. 3.**

**Figure S5:**
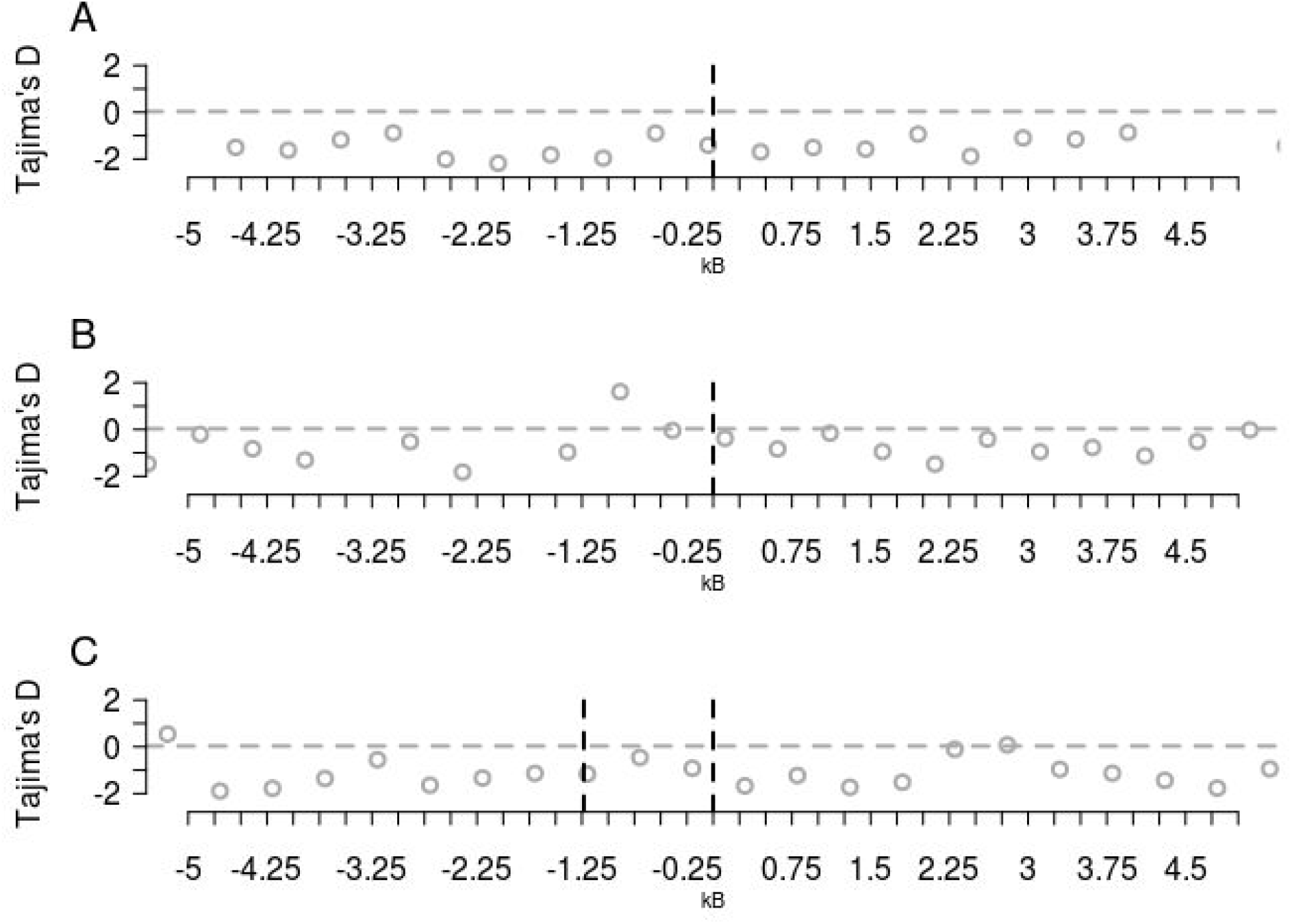
Tajima’s D at putatively neutral sites in 500 bp windows around coexpression-eQTLs. Gray horizontal dotted lines show 95% cutoffs of the observed distribution of likelihoods and black vertical lines show the locations of eQTLs.. Each panel corresponds to the eQTL shown in **Fig. 1.**

**Figure S6:**
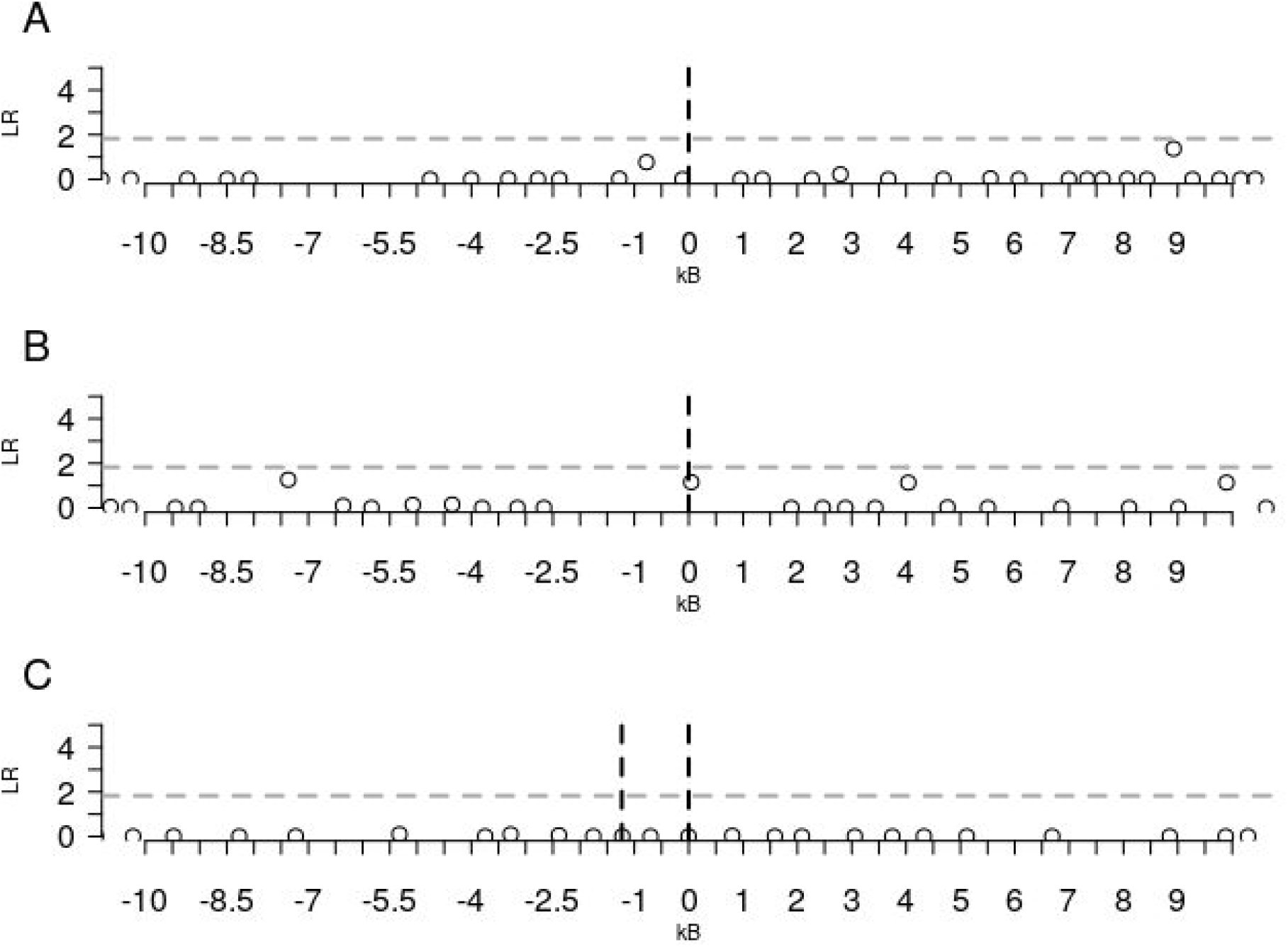
SweeD sweep likelihoods in windows around coexpression-eQTLs. Gray horizontal dotted lines show 95% cutoffs of the observed distribution of likelihoods and black vertical lines show the locations of eQTLs. Each panel corresponds to the eQTL shown in **Fig. 1.**

